# Differential tracking of linguistic vs. mental state content in naturalistic stimuli by language and Theory of Mind (ToM) brain networks

**DOI:** 10.1101/2021.04.28.441724

**Authors:** Alexander M. Paunov, Idan A. Blank, Olessia Jouravlev, Zachary Mineroff, Jeanne Gallée, Evelina Fedorenko

**Affiliations:** Department of Brain and Cognitive Sciences, MIT, Cambridge MA 02139; McGovern Institute for Brain Research, MIT, Cambridge MA 02139; Department of Psychology, UCLA, Los Angeles CA 90095; Institute for Cognitive Science, Carleton University, Ottawa, Canada K1S 5B6; Program in Speech and Hearing Biosciences and Technology, Harvard University, Boston, MA, 02115

**Keywords:** language, theory of mind, social cognition, intersubject correlations, naturalistic fMRI

## Abstract

Language and social cognition, especially the ability to reason about mental states, known as Theory of Mind (ToM), are deeply related in development and everyday use. However, whether these cognitive faculties rely on distinct, overlapping, or the same mechanisms remains debated. Some evidence suggests that, by adulthood, language and ToM draw on largely distinct—though plausibly interacting—cortical networks. However, the broad topography of these networks is similar, and some have emphasized the importance of social content / communicative intent in the linguistic signal for eliciting responses in the language areas. Here, we combine the power of individual-subjects functional localization with the naturalistic-cognition inter-subject correlation approach to illuminate the language-ToM relationship. Using fMRI, we recorded neural activity as participants (n=43) listened to stories and dialogs with mental state content (+linguistic, +ToM), viewed silent animations and live action films with mental state content but no language (-linguistic, +ToM), or listened to an expository text (+linguistic, -ToM). The ToM network robustly tracked stimuli rich in mental state information regardless of whether mental states were conveyed linguistically or non-linguistically, while tracking a +linguistic/-ToM stimulus only weakly. In contrast, the language network tracked linguistic stimuli more strongly than a) non-linguistic stimuli, and than b) the ToM network, and showed reliable tracking even for the linguistic condition devoid of mental state content. These findings suggest that in spite of their indisputably close links, language and ToM dissociate robustly in their neural substrates—and thus plausibly cognitive mechanisms—including during the processing of rich naturalistic materials.

## 1. Introduction

Language and social cognition, especially the ability to reason about mental states, known as Theory of Mind (ToM), are deeply related in human development, everyday use, and possibly evolution. After all, language use is a communicative behavior, which is a kind of cooperative behavior, and cooperative behaviors are, in turn, a kind of social behavior (e.g., Grice, 1968, 1975, Sperber & Wilson, 1986). Construed this way, language can hardly be encapsulated from social cognition (cf. Fodor, 1983); it is subsumed *within* social cognition. Interpreting linguistic signals bears key parallels to the interpretation of other intentional behaviors (e.g., Grice, 1975; Sperber & Wilson, 1986). Communicative utterances, like other behaviors, are assumed to have goals, and conversation partners are assumed to pursue these goals rationally. Furthermore, everyday discourse appears to be dominated by information about other people (e.g., Dunbar, Marriott & Duncan, 1997), and the need to keep track of others’ social record has been proposed as a key driver of language evolution (e.g., Dunbar, 2004; Nowak & Sigmund, 2005; Sommerfield, Krambeck & Milinski, 2008; Nowak & Highfield, 2011). Lastly, evidence of others’ mental states conveyed through language is arguably richer and certainly more direct / less ambiguous than what can be inferred from non-linguistic intentional behavior alone: trying to infer the beliefs guiding someone’s actions can be obviated by their telling you what those beliefs are.

Yet in the relatively brief history of cognitive neuroscience, the study of language and of social cognition have, for the most part, proceeded independently. The two cognitive domains have been treated, often implicitly, as components of a “nearly decomposable system” (Simon, 1962)—a system in which interactions between the component subsystems (language and social cognition here) are sufficiently less common than interactions among each subsystem’s parts to warrant a “divide and conquer” strategy (Saxe, Brett, & Kanwisher, 2006). On such a strategy, each subsystem can first be investigated on its own, leaving questions of potential dependencies between subsystems for a later stage. The application of this strategy to language and social cognition has been remarkably successful, and even this later stage is now coming within reach. But is near-decomposability a true property of the relationship between language and social cognition, or merely a simplifying assumption? It is difficult to say because most studies to date have deliberately aimed to isolate one or the other component subsystem rather than probe their relationship (cf. Deen et al., 2015; Paunov et al., 2019; Braga et al., 2020). So we know that language and social cognition *can* dissociate, under appropriate experimental conditions. But do they, in fact, dissociate in everyday, naturalistic cognition? This is the question we tackle in the current study.

Before we proceed to describe the current study, we first describe two ways in which language and ToM are related in adult cognition and clarify our research aims with respect to these. Then we review what is currently known about the neural bases of linguistic processing and of Theory of Mind.

### 1.1. The role of ToM in inferring communicative intent vs. processing linguistic content

Two aspects of the relationship between language and social cognition can be identified in the context of communication: i) the role of mental state inference in language comprehension generally, whether or not the content concerns mental states; and ii) the use of language to express information about the mental states of agents, either directly or through descriptions of physical events, which prompt mental state attribution (e.g., Fletcher et al., 1995; Gallagher et al., 2000; Saxe & Kanwisher, 2003; Saxe, Schulz, & Jiang, 2006; Jacoby et al., 2016). In the present study, we focus primarily on the second of these aspects.

The first aspect of the language-ToM relationship is generally subsumed in the broad category of *pragmatics,* which is traditionally concerned with how communicative intent—a form of ToM inference—guides linguistic interpretation (e.g., Grice, 1957, 1968, 1975). A key starting point of pragmatics is that the intended meaning of utterances is vastly underdetermined by linguistic inputs (Wittgenstein, 1953; Sperber & Wilson, 1986), and ToM must be recruited to draw inferences about what a speaker intends to convey, based on evidence contained in the speaker’s utterances and the extended context (linguistic, visual, social, etc.). However, delimiting the scope of pragmatic inference is a long-standing challenge, raising the question of whether it is possible to draw a boundary between decoded (literal) and inferred meaning, that is, between semantics and pragmatics (e.g., Jackendoff, 2009).

The lack of clearly defined boundaries for the construct of pragmatics poses empirical challenges. For one, pragmatic inference need not be limited to linguistic communication: it can equally be present in other forms of cooperative information transfer between individuals, such as the interpretation of communicative gestures or, more relevant in the present context, understanding non-linguistic ‘stories’ such as non-verbal animated or live-action films. At the same time, within the narrower context of linguistic communication, it is implausible that all forms of context-based inference of meaning recruit mental state reasoning. Phenomena that require context-based inferences include not just the paradigmatic instances of non-literal meaning, such as irony, indirect requests, hyperbole, and other ‘conversational implicatures’ (Grice, 1975), but also relatively ‘low-level’ and ubiquitous phenomena such as pronoun resolution, or lexical and syntactic disambiguation.

Further, different situational contexts plausibly differ in the nature and extent of their social-cognitive demands. In **Figure 1**, we schematically illustrate three contexts for language processing: reading or listening to narratives or expository texts, reading or listening to a conversation, and directly participating in face-to-face conversation. For each, we outline the ToM cognitive demands associated with, on the one hand, a) understanding the *communicative intent* of the person generating the linguistic output, and, on the other, b) the processing of the *content* of the linguistic materials. Past work has investigated both kinds of ToM demands. For example, a number of studies have manipulated the difficulty of inferring the communicative intent of a speaker by examining the kinds of paradigmatic cases of non-literal language mentioned above, from irony (e.g., Spotorno et al., 2012), to indirect speech (e.g., Feng et al, 2017; van Ackeren et al., 2012) and other forms of conversational implicature (e.g., Feng, Yu, & Zhou, 2021; Jang et al., 2013; see Hagoort & Levinson, 2014 for review). These studies have reported stronger responses in the ToM network for the critical, non-literal stimuli compared to literal controls. Other studies have varied the amount of mental state content in verbal vignettes or stories (e.g., Ferstl & von Cramon, 2002; Saxe & Powell, 2006) and reported stronger activity in the ToM network for materials rich in mental state content (although even expository connected texts appear to engender responses in the ToM network relative to sets of unconnected sentences; Jacoby et al., 2020). So, both kinds of demands appear capable of recruiting mental state reasoning.

**Figure 1.**
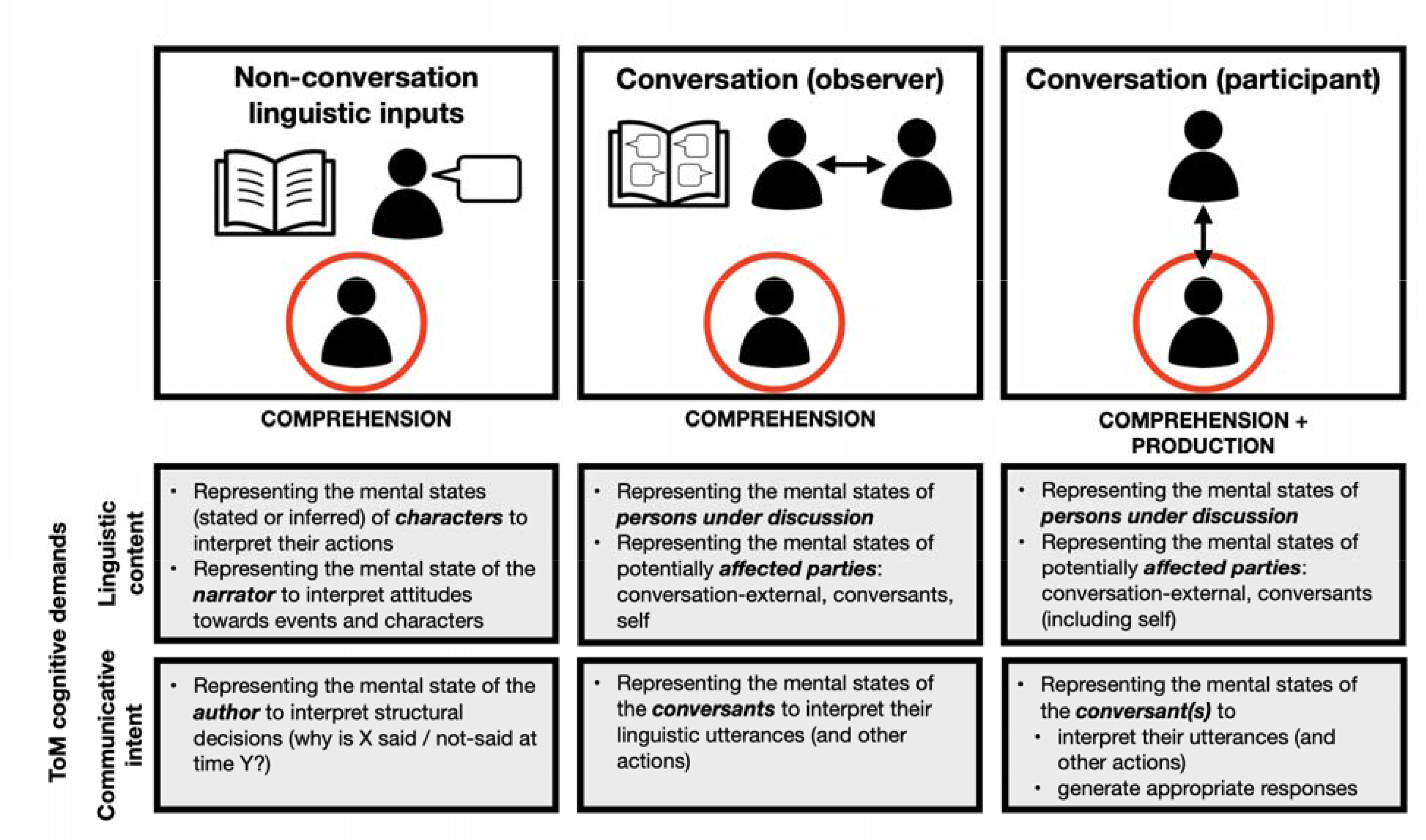
Cognitive demands on ToM processing across different contexts of discourse-level linguistic processing with respect to understanding the communicative intent of the producer, and the linguistic content of the materials. The comprehender is circled in red. Whereas specific demands differ across contexts, any form of language use arguably involves attribution of communicative intent (pragmatics) in the service of comprehension and / or production.

In the current study, we focus on the presence or absence of ToM content in rich naturalistic materials—in line with current emphasis in the field on the importance of going beyond carefully controlled experimental materials in the testing of hypotheses about human cognitive architecture (e.g., Blank, Kanwisher & Fedorenko, 2014; Huth et al., 2016; Hasson et al, 2018)— and evaluate the evidence for *differential recruitment* of language and ToM cortical regions across these materials. Establishing robustly dissociable neural substrates for language understanding and mental state inference is an important step toward bringing neural evidence to bear on the differentiation of pragmatics into distinct sub-domains of context-based inference, which may differentially entail mental state attribution (e.g., Sperber & Wilson, 2002; Hagoort & Levinson, 2014; Bosco, Tirassa, & Gabbatore, 2018).

### 1.2. Neural bases of language and Theory of Mind

#### Language network

Language processing engages a network of left-lateralized brain regions in lateral frontal and temporal cortex. These regions support lexico-semantic processing (word meanings) and combinatorial morpho-syntactic and semantic processing (e.g., Fedorenko et al., 2010, 2012, 2016, 2020; Bautista & Wilson, 2016; Blank et al., 2016; Anderson et al., 2018; Reddy & Wehbe, 2020). Responses to linguistic stimuli generalize across different materials, tasks, modalities of presentation (reading vs. listening), languages, and developmental trajectories (e.g., Fedorenko et al., 2010; Bedny et al., 2011; Fedorenko, 2014; Scott et al., 2017; Diachek, Blank, Siegelman et al., 2020; Ayyash, Malik-Moraleda et al., in prep.; Ivanova et al., in prep.). In contrast to these robust and consistent responses to linguistic stimuli, these regions do not respond to a wide range of non-linguistic cognitive processes, including arithmetic processing, music perception, executive function tasks, action or gesture observation, non-verbal social-cognitive tasks, and the processing of computer code (e.g., Fedorenko et al., 2011; Monti et al., 2012; Pritchett et al., 2018; Almaric & Dehaene, 2018; Paunov, 2018; Jouravlev et al., 2019; Ivanova et al., 2020; Shain, Paunov et al., in prep.; see Fedorenko & Varley, 2016 for a review).

#### ToM network

Attribution of mental states, such as beliefs, desires, and intentions, engages a network of brain regions in bilateral temporo-parietal cortex and anterior and posterior regions along the cortical midline. These responses generalize across the type of mental state, its specific content or format (linguistic vs. pictorial), and the source of evidence for it (e.g., Fletcher et al., 1995; Castelli et al., 2000; Gallagher et al., 2000; Vogeley et al., 2001; Ruby & Decety, 2003; Saxe & Kanwisher, 2003; Saxe, Schulz, & Jiang, 2006; Kandylaki et al., 2015; Jacoby et al., 2016; see Koster-Hale & Saxe, 2013 for review) as well as across populations with diverse cultural backgrounds or developmental experiences (e.g., Callaghan et al., 2005; Sabbagh et al., 2006; Shahaeian et al., 2011; Koster-Hale, Bedny, & Saxe, 2014; Richardson et al., 2018, 2020). However, by adulthood, these regions, and especially the most selective component of the ToM network, the right temporo-parietal junction (RTPJ), do not respond to social stimuli, like faces, voices, or biological motion (e.g., Deen et al., 2015), to general executive demands (e.g., Saxe, Schulz, & Jiang, 2006), to physical or broadly social attributes of agents, or to attribution of bodily sensations of pain and hunger (e.g., Saxe & Powell, 2006; Bruneau, Pluta & Saxe, 2012; Jacoby et al., 2016).

Together, these two lines of work support the notion that language and ToM are distinct cognitive faculties, each mapping onto functionally specialized neural structures. Investigations of developmental and acquired disorders have provided convergent support for the dissociability of language and ToM. Individuals with even severe aphasia appear to retain the capacity for mental state reasoning as long as non-verbal materials are used (e.g., Varley & Siegal, 2000; Varley, Siegal & Want, 2001; Apperly et al., 2006; Willems et al., 2011; see Fedorenko & Varley, 2016 for review). And at least some individuals with social, ToM-related impairments (e.g., some individuals with autism spectrum disorders) show preservation of lexical and syntactic linguistic abilities (e.g., Frith & Happe, 1994; Wilkinson, 1998; Tager-Flusberg, Paul & Lord, 2005; Tager-Flusberg, 2006). But as alluded to above, most of this evidence has come from traditional, task-based experimental paradigms, and such paradigms are well-suited to the divide-and-conquer enterprise, but may overestimate the degree of separation. Indeed, Paunov et al. (2019) recently examined inter-regional functional correlations within and between the language and ToM networks during naturalistic cognition paradigms, like story comprehension, and found that in spite of their dissociability (stronger within-than between-network correlations), the language and ToM networks also showed a significant amount of synchronization in their neural activity. These results point to some degree of functional integration between the networks. Furthermore, several prior studies have argued for the importance of social content / communicative intent in the linguistic signal for eliciting responses in the language areas (e.g., Mellem et al., 2016; Redcay et al., 2016). Here, we adopt an approach that relies on the use of rich naturalistic stimuli but allows us to not only examine the structure of and the relationship between the networks, but to also ask what stimulus features each network “tracks” through the inter-subject correlation approach (e.g., Hasson et al., 2004, 2008).

## 2. Materials and Methods

### 2.1. General approach

Following Blank & Fedorenko (2017, 2020), we combine two powerful methodologies, which have previously been productively applied separately in the domains of language and social cognition. In particular, we use *functional localization* (e.g., Brett et al., 2002; Saxe et al., 2006; Fedorenko et al., 2010) to identify the two networks of interest in individual subjects, and *inter-subject correlations* (ISCs; e.g., Hasson et al., 2004, 2008) to examine the degree to which these networks ‘track’ different stimulus features during the processing of rich naturalistic materials. Here, we highlight key strengths of each approach and consider their synergistic advantages in the context of our research goals.

Naturalistic paradigms have become a crucial complement to traditional, task-based studies in cognitive neuroscience. The obvious advantage of naturalistic paradigms is their high ecological validity: by giving up a measure of experimental control, it becomes possible to study cognition “in the wild” (Blank & Fedorenko, 2017; see e.g., Sonsukare et al., 2019 and Nastase et al., 2020 for general discussions). In particular, one can examine how coherent and structured mental representations are extracted from rich and noisy perceptual inputs, which is what happens in everyday cognition. This is in contrast to artificially isolating various features of these perceptual inputs, as is typically done in constrained experimental tasks. Naturalistic paradigms have been argued to elicit more reliable responses compared to traditional, task-based paradigms (Hasson et al., 2010), perhaps because they are generally more engaging, and can enable discoveries of functional relationships among brain regions and networks that are altogether missed in more constrained settings (e.g., Gallivan, Cavina-Pratesi & Culham, 2009). Another advantage of naturalistic paradigms is their hypothesis-free nature. Through the use of naturalistic materials, researchers impose minimal design constraints to investigate the domain of interest in a manner that is maximally unbiased by prior theoretical assumptions. In effect, they are letting the data speak for itself.

However, naturalistic paradigms also come with an inherent analytic challenge: how do we make sense of data acquired without the typical constraints of standard hypothesis-driven modeling approaches? Hasson et al. (2004) pioneered an approach to tackle this challenge, known as the inter-subject correlation (ISC) approach (see Hasson et al., 2008, for an overview), which we adopt in the current study. The key insight behind the ISC approach is that we can model any given participant’s fMRI signal time series using another participant’s or other participants’ time series: if a voxel, brain region, or brain network ‘tracks’ features of the stimulus during which the time series were obtained, then fMRI signal fluctuations will be stimulus-locked, resulting in similar time courses across participants (i.e., high inter-subject correlations).

ISCs have been used in several studies of narrative comprehension (e.g., Wilson et al., 2007; Lerner et al., 2011; Honey et al., 2012; Regev et al, 2013; Silber et al., 2014; Schmälzle et al., 2015; Blank & Fedorenko, 2017, 2020), and whole-brain voxel-wise analyses have revealed high inter-subject correlations across large swaths of cortex that resemble the union of the language and ToM networks. On their own, these results might be taken as *prima facie* evidence for non-dissociability of language and ToM, given that the two networks appear to be jointly recruited.

And insofar as the mental processes recruited in narrative comprehension recapitulate those used in everyday abstract cognition—an assumption that, we take it, partially justifies the interest in narratives in cognitive science and neuroscience (e.g., Finlayson & Winston, 2011; Willems et al., 2020)—the results may be taken to suggest the non-dissociability of language and ToM more generally.

However, it is difficult to draw inferences from these studies about the *relative* contributions of the language and ToM networks to narrative comprehension for two reasons. First, in whole-brain analyses, ISCs are computed on a voxel-wise basis: individual brains are normalized to a stereotaxic template, and one-to-one voxel correspondence across individuals is then assumed in computing the ISCs. This approach is problematic because i) inter-individual variability is well established in the high-level association cortex (e.g., Fischl et al., 2008; Frost & Goebel, 2012; Tahmasebi et al., 2012), so any given voxel may belong to functionally distinct regions across participants; and ii) there is no independent criterion based on which an anatomical location can be interpreted as belonging to the language vs. the ToM network, thus necessitating reliance on the fallacious “reverse inference” (Poldrack, 2006) to interpret the resulting topography (see Fedorenko, 2021, for discussion). And second, traditional whole-brain analyses typically include all voxels that showed significant (above baseline) ISCs, thus potentially obscuring large differences in effect sizes (cf. Blank & Fedorenko, 2017; see Chen, Taylor & Cox, 2017, for a general discussion of the importance of considering effect sizes in interpreting fMRI findings). Combining the ISC approach with individual-participant functional localization enables us to identify and directly compare the networks of interest (including with respect to effect sizes), as well as to relate the findings straightforwardly to the prior literature on the language and ToM networks. In the current study, we therefore identified the language and ToM networks using well-established functional localizers (Saxe & Kanwisher, 2003; Fedorenko et al., 2010), and then examined the degree of inter-subject synchronization in those regions during the processing of diverse naturalistic linguistic and non-linguistic conditions varying in the presence of mental state content. If the language and ToM networks are dissociable during naturalistic cognition, we would expect the language regions to track linguistic stimuli, including those that lack mental state content, and the ToM regions to track stimuli that have mental state content, including both linguistic and non-linguistic ones.

### 2.2. Overall experimental design and statistical analyses

Our overall design and analytic strategy were as follows: participant-specific regions that responded more strongly during the reading of sentences compared with lists of nonwords were defined as regions of interest comprising the language network. Similarly, regions that responded more strongly to stories about others’ beliefs vs. stories about physical reality were defined as regions of interest comprising the ToM network (see ***Section 2.4*** for details). Whereas the precise anatomical locations of these regions were allowed to vary across participants, their overall topography was constrained by independently derived criteria to establish functional correspondence across brain regions of different participants (e.g., Fedorenko et al., 2010; Julian et al., 2012).

Activity in these two sets of brain regions was recorded with fMRI while participants listened to or watched a series of naturalistic stimuli, as detailed below. For each region in each network, our critical dependent variable was the strength of the correlation between each participant’s time series and the average time series from the rest of the participants. The group-averaged ISC in each region was tested for significance via a permutation test of the time series data. For our critical analysis, all individual ISC values were modeled using a linear mixed-effects regression with participant, brain region, and stimulus (what we call ‘condition’ below) as random effects.

### 2.3. Participants

Forty-seven native English speakers (age 19-48, M = 24.5, SD = 5.08; 30 female) from MIT and the surrounding Boston community participated for payment. Forty participants were right-handed, as determined by the Edinburgh handedness inventory (Oldfield 1971) or by self-report (n=1). All seven left-handed participants showed typical left lateralization in the language localizer task described below (see Willems et al., 2014, for arguments to include left handers in cognitive neuroscience research). Four participants were excluded from the analyses due to poor quality of the localizer data (2 for ToM localizer, 1 for language localizer, and 1 for both), with the exclusion criterion defined as fewer than 100 suprathreshold voxels (at the p *<* 0.001 uncorrected whole-brain threshold) across the respective network’s masks (see below), bringing the number of participants included in the critical analyses to 43. All participants gave informed written consent in accordance with the requirements of MIT’s Committee on the Use of Humans as Experimental Subjects (COUHES).

### 2.4. Stimuli and procedure

Each participant completed a language localizer, a ToM localizer, a localizer for the domain-general Multiple Demand (MD) system (used in a replication analysis, as described below), and a subset, or all, of the critical naturalistic stimuli (‘conditions’) (between 1 and 7) due to scan duration constraints (18 participants completed all 7 conditions of interest, 1 participant completed 6 conditions, 10 participants completed 5 conditions, 1 participant completed 4 conditions, 1 participant completed 3 conditions, 7 participants completed 2 conditions, and 5 participants completed 1 condition). Each condition was presented to between 28 and 32 participants (see **Table 1**). Each stimulus, lasting ∼5-7 minutes (see **Table 1** for precise durations), was preceded and followed by 16 s of fixation. Finally, ten participants performed a resting state scan, used in one of the reality-check analyses, as described below. For the language localizer, 36/43 participants completed it in the same session as the critical conditions, the remaining 7 participants completed it in an earlier session. Similarly, for the ToM and MD localizers, 37/43 participants completed them in the same session as the critical conditions, the remaining 6 participants completed them in an earlier session. We will now describe the localizers and the critical experiment in more detail.

**Table 1.**
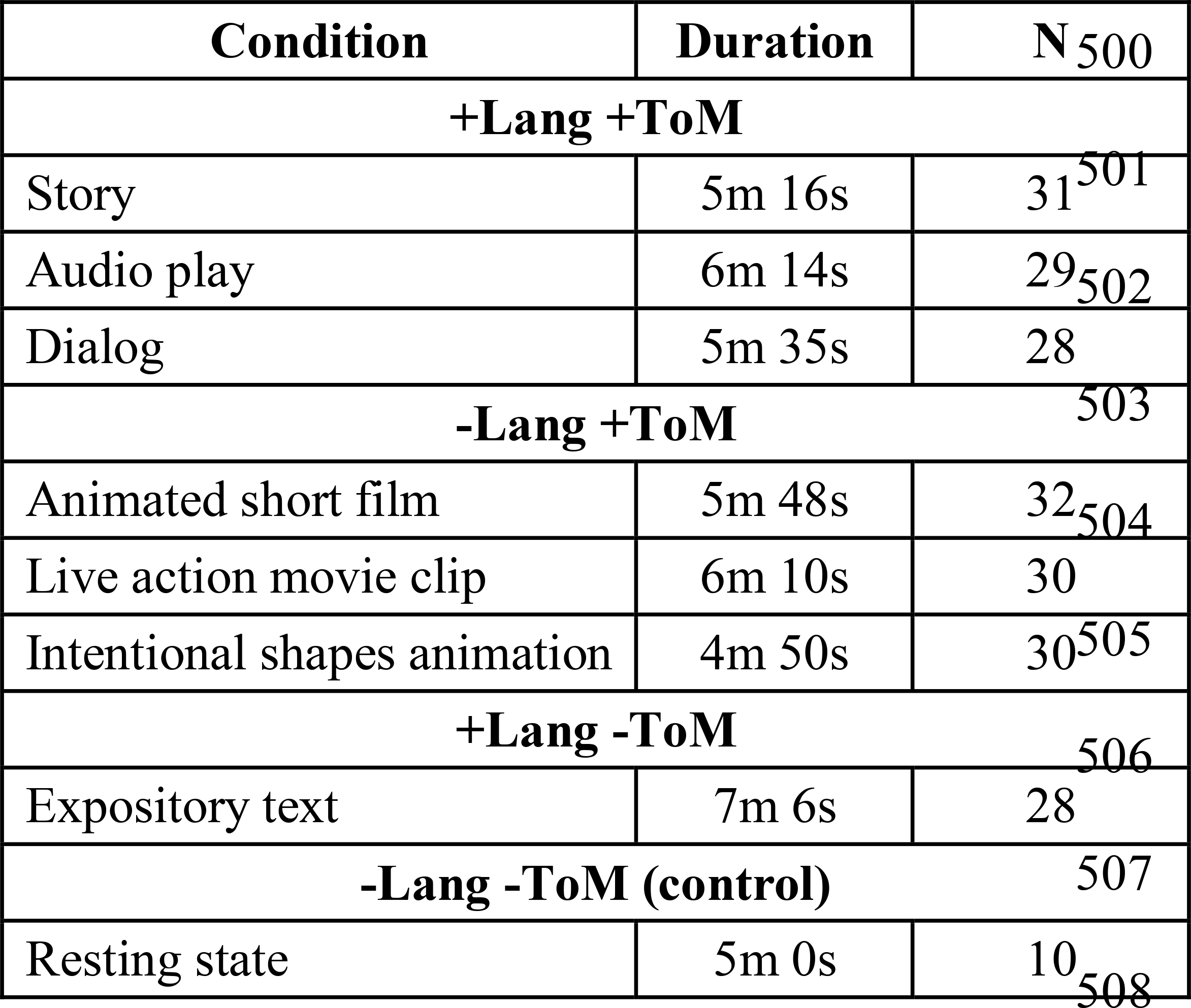
Naturalistic conditions in each of the three condition types of interest and a reality-check condition (resting state), including durations, and number of participants per condition. Durations include 16 s fixations at the beginning and end of the scan (32 s total).

| Condition | Duration | N |
| --- | --- | --- |
| **+Lang +ToM** |  |  |
| Story | 5m 16s | 31<sup>501</sup> |
| Audio play | 6m 14s | 29<sup>502</sup> |
| Dialog | 5m 35s | 28 |
| **-Lang +ToM** |  |  |
| Animated short film | 5m 48s | 32<sup>504</sup> |
| Live action movie clip | 6m 10s | 30 |
| Intentional shapes animation | 4m 50s | 30<sup>505</sup> |
| **+Lang -ToM** |  |  |
| Expository text | 7m 6s | 28 |
| **-Lang -ToM (control)** |  |  |
| Resting state | 5m 0s | 10<sup>508</sup> |

#### Language localizer task

The task used to localize the language network is described in detail in Fedorenko et al. (2010) and targets brain regions that support high-level language processing. Briefly, we used a reading task contrasting sentences and lists of unconnected, pronounceable nonwords (**Figure 2**) in a blocked design with a counterbalanced condition order across runs. Stimuli were presented one word / nonword at a time. Participants read the materials passively (we included a button-pressing task at the end of each trial, to help participants remain alert). As discussed in the introduction, this localizer is robust to task manipulations (e.g., Fedorenko et al., 2010; Scott et al., 2017; Ivanova et al., in prep.). Moreover, this localizer identifies the same regions that are localized with a broader contrast, between listening to natural speech and its acoustically-degraded version (Scott et al., 2017; Ayyash, Malik-Moraleda et al., in prep.). All participants completed two runs, each lasting 358 s and consisting of 8 blocks per condition and 5 fixation blocks. (A version of this localizer is available from http://evlab.mit.edu/funcloc/.)

**Figure 2.**
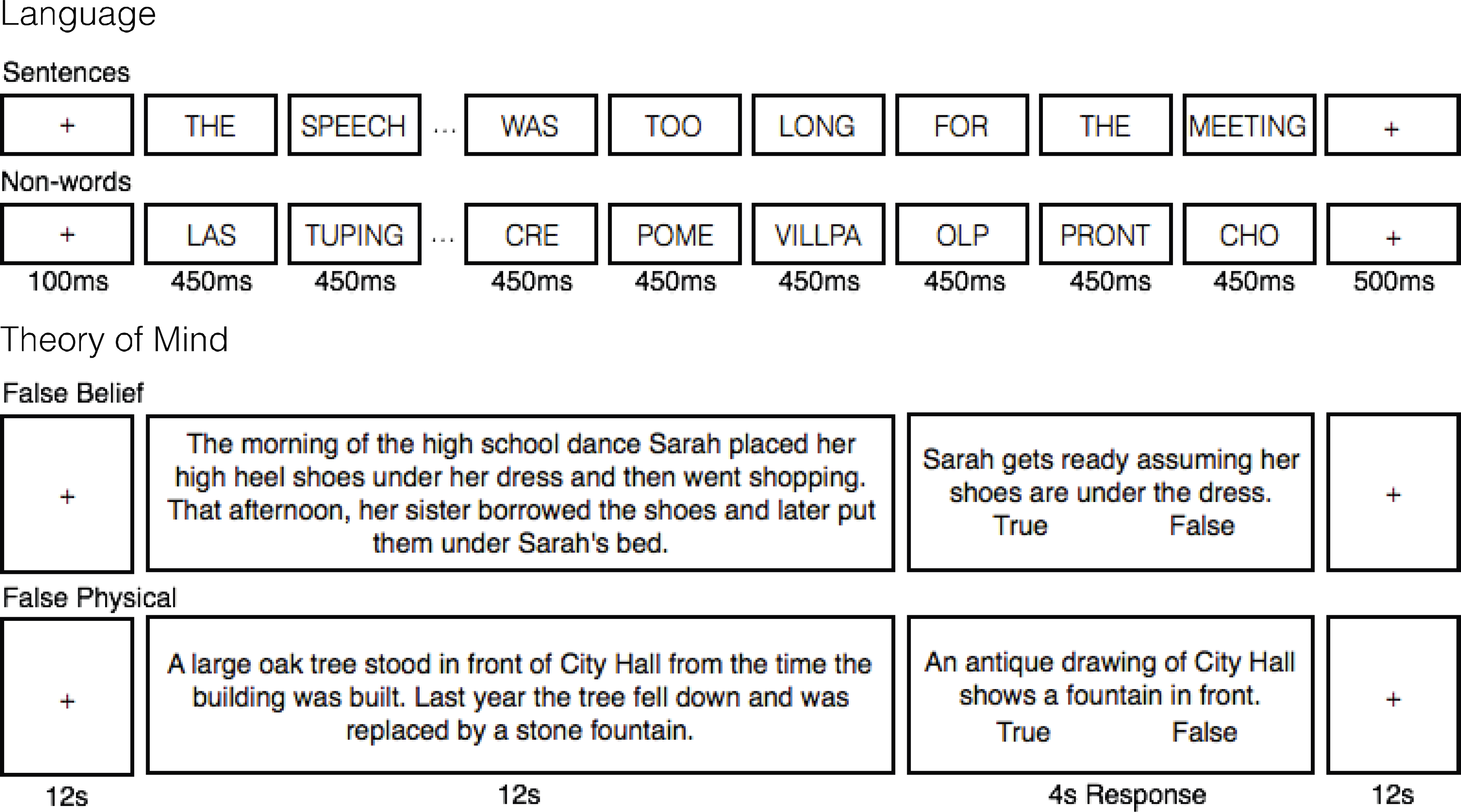
Sample trials from the functional localizer paradigms. **Language**: reading of sentences was contrasted with reading of sequences of pronounceable non-words (Fedorenko et al., 2010). **ToM**: reading of vignettes about false mental states was contrasted with reading of vignettes about false physical states, each followed by a true/false statement (Saxe & Kanwisher, 2003).

#### ToM localizer task

The task used to localize the ToM network is described in detail in Saxe & Kanwisher (2003) and targets brain regions that support reasoning about others’ mental states. Briefly, the task was based on the classic false belief paradigm (Wimmer and Perner, 1983), and contrasted verbal vignettes about false beliefs (e.g., a protagonist has a false belief about an object’s location; the critical condition) and linguistically matched vignettes about false physical states (physical representations depicting outdated scenes, e.g., a photograph showing an object that has since been removed; the control condition) (**Figure 2**) in a long-event-related design with a counterbalanced order across runs (when multiple runs were administered). Stimuli were presented one at a time. Participants read these vignettes and answered a true / false comprehension question after each one. Forty-one participants completed two runs and two completed one run due to time limitations, each lasting 262 s and consisting of 5 vignettes per condition. (A version of this localizer is available from http://saxelab.mit.edu/use-our-efficient-false-belief-localizer.)

#### An alternative, non-verbal ToM localizer task

One of the naturalistic conditions in this study (“Partly Cloudy”, an animated short film; see next section) has been previously used as a non-verbal ToM localizer (Jacoby et al., 2016; see also Richardson et al., 2018). To that end, it has been coded into ‘mental’, ‘physical’, ‘social’, and ‘pain’ segments, and the regions defined by the mental > pain contrast have been validated against the traditional ToM localizer described above (see Jacoby et al., 2016 for details). Examples of mental content include a character falsely believing they have been abandoned by a companion and a character observing others interacting happily after experiencing pain (4 events, 44 sec total). Following a reviewer’s suggestion, we used this localizer as an alternative ToM localizer in some of the analyses. We used the mental > physical contrast rather than mental > pain to maintain conceptual similarity with the verbal localizer’s false belief > false physical contrast. The activations obtained with the two different control conditions were qualitatively similar. Examples of physical content include a wide shot of clouds and birds flying (3 events, 22 sec total). The main goal was to ensure that the language-ToM dissociation is not due to an overly narrow definition of Theory of Mind in terms of false beliefs, implicit in the use of this particular type of mental state attribution in the standard, verbal localizer. Notably, many previous studies have shown that the ToM network defined with the false belief localizer responds to a wide range of mental state content besides (false) beliefs, including intentions, sources of evidence about others’ minds, emotional pain, and the ‘minds’ of group agents (e.g., Young & Saxe, 2008; Bruneau, Pluta & Saxe, 2012; Koster-Hale, Bedny & Saxe, 2014; Jenkins et al., 2014). Nevertheless, analyses that use the non-verbal ToM localizer should confirm that the results of the present study generalize beyond a particular way of localizing the ToM network.

*Multiple Demand (MD) localizer task* (used in a replication analysis, as described below). The task used to localize the MD network is described in detail in Fedorenko et al. (2011) and targets brain regions that support goal-directed effortful behaviors (e.g., Duncan, 2010, 2013). Briefly, we used a spatial working-memory task contrasting a harder version with an easier version (**Supplemental Figure A1**) in a blocked design with a counterbalanced condition order across runs (when multiple runs were administered). On each trial, participants saw a 3x4 grid and kept track of eight (hard version) or four (easy version) randomly generated locations that were sequentially flashed two at a time or one at a time, respectively. Then, participants indicated their memory for these locations in a two-alternative, forced-choice paradigm via a button press, and received feedback. Of the thirty-two participants included in the replication analysis (i.e., non-overlapping with those used in the original study in Blank & Fedorenko, 2017), twenty-three participants completed two runs of the localizer, and nine completed one run, each lasting 448 s and consisting of 6 blocks per condition and 4 fixation blocks.

#### The critical naturalistic task

In the main experiment, each participant listened to (over scanner-safe Sensimetrics headphones) and/or watched a set of naturalistic stimuli (varying between 4 min 50 s and 7 min 6 s in duration). Four of the conditions used linguistic materials: i) a story (“Elvis” from the Natural Stories corpus; Futrell et al., 2020), ii) an audio play (a segment from an HBO miniseries, “Angels in America”, from Chapter 1: “Bad News”, https://www.imdb.com/title/tt0318997/, audio only), iii) a naturalistic dialog—a casual unscripted conversation between two female friends (recorded by JG), and iv) a non-narrative expository text (a text about trees adapted from Wikipedia; https://en.wikipedia.org/wiki/Tree) (recorded by JG). The first three of the linguistic conditions were rich in mental state content; the fourth was meaningful naturalistic discourse with little/no mental state content (see below for additional discussion). The three remaining conditions were videos with no linguistic content: i) an animated short film (“Partly Cloudy” by Pixar), ii) a clip from a live action film (“Mr. Bean”, https://www.youtube.com/watch?v=bhg-ZZ6WA), and iii) a custom-created Heider and Simmel style animation (Heider & Simmel, 1944) consisting of simple geometric shapes moving in ways as to suggest intentional interactions designed to tell a story (e.g., a shape gets locked up inside a space, another shape goes on a quest to get help to release it, etc.). All three non-linguistic conditions were rich in mental state content. (Five additional conditions—included in some participants for another study—are of no interest to the current study.) All the materials are available on OSF (except in cases where copyright issues will prevent us from doing so): https://osf.io/prghx/. In the resting state scan, used for one of our reality-check analyses, as described below, participants were instructed to close their eyes and let their mind wander but to remain awake while resting in the scanner for 5 min (the scanner lights were dimmed and the projector was turned off).

It is important to note that although we classify these naturalistic conditions into ‘types’ in a binary way (i.e., either involving ToM or not, and either involving language or not), this should not be taken to suggest that there cannot be gradation within each category. Indeed, given the richness of the stimuli, there almost certainly is, at least for the ToM dimension. However, we do not pursue this question further given the small number of stimuli within each category, and their complexity. Another challenge is coding the materials, especially for mental state content. In particular, linguistically mediated mental state attribution often proceeds not from explicit mentions of mental states but from action descriptions, and in non-verbal settings, ToM attribution proceeds exclusively from action observation. As a result, it is difficult to specify, especially in complex naturalistic materials, when mental state attribution is prompted.

Conversely, ToM vocabulary need not lead to stronger mental state attribution (e.g., “Alan thinks that this is a nice house” *vs.* “Alan says that this is a nice house”; presumably, a mental state is ascribed to Alan in both cases). Thus, we do not attempt to quantify degrees of mental state attribution beyond the overall presence or (near-) absence of mental state content.

Another important point to acknowledge is that there may be a certain degree of “contamination” across categories. Specifically, as discussed in the Introduction, the ‘-ToM’ condition (expository text) plausibly involves some degree of pragmatic inference, and such texts have previously been shown to elicit responses in parts of the ToM network (e.g., Jacoby & Fedorenko, 2020; Ferstl & von Cramon, 2002). In the case of language, even though binary classification is relatively straightforward (linguistic input is either present or not), one might argue that the language network may nevertheless be recruited to some extent owing to the communicative nature or rich semantics of the non-linguistic stimuli. Although prior work suggests that non-verbal communication does not recruit the language network (e.g., Deen et al., 2015; Pritchett et al., 2018; Jouravlev et al., 2019), some studies have found the language network is activated during the processing of visual event semantics (e.g., Ivanova et al., 2020). Critically for present purposes, however, to the extent that there is ‘contamination’ in either direction, it should work *against* finding a language-ToM dissociation; i.e., our results might underestimate the true degree of dissociation. The rich and graded nature of the stimuli might help account for some results inconsistent with a *complete* dissociation, and we return to this question in the Discussion.

### 2.5. Data acquisition and preprocessing

#### Data acquisition

Whole-brain structural and functional data were collected on a whole-body 3 Tesla Siemens Trio scanner with a 32-channel head coil at the Athinoula A. Martinos Imaging Center at the McGovern Institute for Brain Research at MIT. T1-weighted structural images were collected in 176 axial slices with 1 mm isotropic voxels [repetition time (TR) = 2530 ms; echo time (TE) = 3.48 ms]. Functional, blood oxygenation level-dependent (BOLD) data were acquired using an EPI sequence with a 90° flip angle and using GRAPPA with an acceleration factor of 2; the following parameters were used: thirty-one 4-mm-thick near-axial slices acquired in an interleaved order (with 10% distance factor), with an in-plane resolution of 2.1 x 2.1 mm, FoV in the phase encoding (A >> P) direction 200 mm and matrix size 96 x 96 mm, TR = 2000 ms and TE = 30 ms. The first 10 s of each run were excluded to allow for steady-state magnetization.

#### Spatial preprocessing

Data preprocessing was performed with SPM12 (using default parameters, unless specified otherwise) and supporting, custom scripts in MATLAB. Preprocessing of anatomical data included normalization into a common space [Montreal Neurological Institute (MNI) template], resampling into 2 mm isotropic voxels, and segmentation into probabilistic maps of the gray matter, white matter (WM), and cerebro-spinal fluid (CSF). Preprocessing of functional data included motion correction (realignment to the mean image using second-degree b-spline interpolation), normalization (estimated for the mean image using trilinear interpolation), resampling into 2 mm isotropic voxels, smoothing with a 4 mm FWHM Gaussian filter and high-pass filtering at 200 s.

#### Temporal preprocessing

Additional preprocessing of data from the story comprehension runs was performed using the CONN toolbox (Whitfield-Gabrieli and Nieto-Castañón, 2012; http://www.nitrc.org/projects/conn) with default parameters, unless specified otherwise. Five temporal principal components of the BOLD signal time courses extracted from the WM were regressed out of each voxel’s time course; signal originating in the CSF was similarly regressed out. Six principal components of the six motion parameters estimated during offline motion correction were also regressed out, as well as their first time derivative. Next, the residual signal was bandpass filtered (0.01 – 0.1 Hz) to preserve only low-frequency signal fluctuations (Cordes et al., 2001). This filtering did not influence the results reported below.

### 2.6. Participant-specific functional localization of the language and ToM (and MD, for a replication analysis) networks

#### Modeling localizer data

For each localizer task, a standard mass univariate analysis was performed in SPM12 whereby a general linear model estimated the effect size of each condition in each experimental run. These effects were each modeled with a boxcar function (representing entire blocks) convolved with the canonical hemodynamic response function. The model also included first-order temporal derivatives of these effects, as well as nuisance regressors representing entire experimental runs, offline-estimated motion parameters, and timepoints classified as outliers (i.e. where the scan-to-scan differences in global BOLD signal are above 5 standard deviations, or where the scan-to-scan motion is above 0.9mm). The obtained weights were then used to compute the functional contrast of interest: for the language localizer, sentences > nonwords, for the ToM localizer false belief > false photo, and for the MD localizer (replication analysis; see ***Section 2.4***), hard > easy spatial working memory.

#### Defining fROIs

Language and ToM (and MD, in the replication analysis) functional regions-of-interest (fROIs) were defined individually for each participant based on functional contrast maps from the localizer experiments (a toolbox for this procedure is available online; https://evlab.mit.edu/funcloc/). These maps were first restricted to include only gray matter voxels by excluding voxels that were more likely to belong to either the WM or the CSF based on SPM’s probabilistic segmentation of the participant’s structural data.

Then, fROIs in the language network were defined using group-constrained, participant-specific localization (Fedorenko et al., 2010). For each participant, the map of the sentences > nonwords contrast was intersected with binary masks that constrained the participant-specific language network to fall within areas where activations for this contrast are relatively likely across the population. These masks are based on a group-level representation of the contrast obtained from a previous sample of 220 participants. We used five such masks in the left-hemisphere, including regions in the mid-to-posterior and anterior temporal lobe, as well as in the middle frontal gyrus, the inferior frontal gyrus, and its orbital part (**Figure 4**). A version of these masks is available online (https://evlab.mit.edu/funcloc/). In each of the resulting 5 masks, a participant-specific language fROI was defined as the top 10% of voxels with the highest contrast values. This top *n*% approach ensures that fROIs can be defined in every participant and that their sizes are the same across participants, allowing for generalizable results (Nieto-Castañon and Fedorenko, 2012).

For the ToM fROIs, we used masks derived from a group-level representation for the false belief > false physical contrast in an independent group of 462 participants (Dufour et al., 2013). These masks included regions in the left and right temporo-parietal junction (L/RTPJ), precuneus / posterior cingulate cortex (L/RPC), and dorsal medial prefrontal cortex (**Figure 4**). A version of these masks is available online (http://saxelab.mit.edu/use-our-theory-mind-group-maps), but the masks were edited as follows: the RSTS (right superior temporal sulcus) mask was excluded, as it covers the entire STS, which is known to show complex functional organization, with reduced ToM selectivity (Deen et al., 2015). The middle- and ventral-medial prefrontal cortex (MMPFC and VMPFC) masks were also excluded to reduce the number of statistical comparisons in per- fROI analyses, but the dorsal MPFC and PC masks were split into left- and right-hemispheres, for a total of 6 masks.

Additionally, for the replication analysis, fROIs in the MD network were defined based on the hard > easy contrast in the spatial working memory task. Here, instead of using binary masks based on group-level functional data, we used anatomical masks (Tzourio-Mazoyer et al., 2002; Fedorenko et al., 2013; Blank et al., 2014; Blank & Fedorenko, 2017). Nine masks were used in each hemisphere, including regions in the middle frontal gyrus and its orbital part, the opercular part of the inferior frontal gyrus, the precentral gyrus, the superior and inferior parts of the parietal lobe, the insula, the supplementary motor area, and the cingulate cortex (**Supplemental Figure A2**). (We note that functional masks derived for the MD network based on 197 participants were largely overlapping with the anatomical masks; we chose to use the anatomical masks to maintain comparability between our functional data and data from previous studies that have used these masks.)

In line with prior studies (e.g., Blank, Fedorenko & Kanwisher, 2014; Blank & Fedorenko, 2017; Paunov, Blank, & Fedorenko, 2019), the resulting fROIs showed small pairwise overlaps within individuals across networks, and overlapping voxels were excluded in fROI definition. In the current sample, the language-MD and ToM-MD overlaps were negligible, with a median overlap of 0 and average percentage overlap of fewer than 3% of voxels, on average across participants, relative to the total size of all fROIs in either network. Similarly, the language-ToM overlaps were small relative to all fROIs in either network (6.3% of voxels, on average across participants, relative to the total number of voxels in language fROIs and 3.2% relative to all ToM fROIs). This overlap was localized entirely to one pair of fROIs: the LPostTemp language fROI and the LTPJ ToM fROI, and was more substantial relative to the total sizes of just these two fROIs: 38.6 voxels, on average across participants, i.e., 13.1% out of 295 total LPostTemp voxels, and 11.6% out of 332 total LTPJ voxels. We therefore repeated all key analyses without excluding these voxels in defining the fROIs. The results of these alternative analyses were qualitatively and statistically similar.

### 2.7. Reality check and replication analyses

Prior to performing our critical analyses, we conducted two reality-check analyses and—in line with increasing emphasis in the field on robust and replicable science (e.g., Poldrack et al., 2017)—an analysis aimed at replicating and extending a previous ISC-based finding from our lab (Blank & Fedorenko, 2017).

#### ISCs in perceptual cortices

Anatomical ROIs were additionally defined in early visual and auditory cortex in all participants. For visual cortex, regions included inferior, middle, and superior occipital cortex bilaterally (six ROIs in total; masks available from http://fmri.wfubmc.edu/software/PickAtlas). For auditory cortex, regions included posteromedial and anterolateral sections of Heschl’s gyrus bilaterally (4 ROIs in total; Morosan et al., 2001; these regions are based on post-mortem histology, and have been used in a number of previous fMRI papers). Signal extraction, ISC estimation, and inferential statistics were performed identically to the critical analyses (see ***Section 2.8***). ISCs from these regions were used in a reality check (see ***Section 3.1***), to ensure a double-dissociation obtains between visually and auditorily presented conditions in these perceptual regions.

#### Resting state ISCs

A subset of ten participants (who completed 1-7 of the critical conditions) completed a resting state scan, which was included to ensure that data acquisition, preprocessing, and modelling procedures do not induce spurious ISCs. To this end, signal extraction, ISC estimation, and inferential statistics were performed identically to the critical analyses (see ***Section 2.8***).

#### Replication analysis: Closer tracking of linguistic input by language regions than by domain-general Multiple Demand (MD) regions

Blank & Fedorenko (2017) reported stronger ISCs during the processing of naturalistic linguistic materials in the language regions, compared to domain-general Multiple Demand (MD) regions. The MD network has been implicated in executive processes and goal-directed behavior (e.g., Duncan, 2010, 2013), including in the domain of language (e.g., see Fedorenko, 2014, for a review; cf. Diachek, Blank, Siegelman et al., 2020; Fedorenko & Shain, under review). We sought to replicate Blank & Fedorenko’s key result in a new set of participants and to extend it to different types of linguistic materials. Specifically, the original study used narratives, including the narrative used in the present study along with three others. We expected the results to generalize to non-narrative linguistic conditions. Ten participants in our dataset (n=6 in the narrative condition) who also participated in the original study were excluded from this analysis. Again, signal extraction, ISC estimation, and inferential statistics were identical to the critical analyses (see ***Section 2.8***).

### 2.8. Critical analyses

#### Computing ISCs

For each participant and fROI, BOLD signal time courses recorded during each naturalistic condition were extracted from each voxel beginning 6 s following the onset of the stimulus (to exclude an initial rise in the hemodynamic response relative to fixation, which could increase ISCs) and averaged across voxels. For each fROI, participant, and condition we computed an ISC value, namely, Pearson’s moment correlation coefficient between the z-scored time course and the corresponding z-scored and averaged time course across the remaining participants (Lerner et al., 2011). ISCs were Fisher-transformed before statistical testing to improve normality (Silver and Dunlap, 1987).

#### Statistical testing

In each fROI, ISCs were then tested for significance against an empirical null distribution based on 1,000 simulated signal time courses that were generated by phase-randomization of the original data (Theiler et al., 1992). Namely, we generated null distributions for individual participants, fit each distribution with a Gaussian, and analytically combined the resulting parameters across participants. The true ISCs, also averaged across participants, were then z-scored relative to these empirical parameters and converted to one-tailed *p*-values.

ISCs were compared across networks and condition types using linear, mixed-effects regressions, implemented in Matlab 2020a. ISCs were modeled with maximal random effects structure appropriate for each analysis (Barr et al., 2013; Baayen et al., 2008), including random intercepts for participants, with random slopes for the effects of interest, and crossed random intercepts for fROI and condition. Hypothesis testing was performed with two-tailed tests over the respective model coefficients, with Satterthwaite approximation for the degrees of freedom.

Further analyses performed within networks across condition types or against the theoretical null distribution (i.e., testing the intercept term), as well as those per fROI within a network or per condition across networks, also always included maximal random effects on the remaining grouping variables. P-values in these analyses are reported following false discovery rate (FDR) correction for multiple comparisons (Benjamini and Yekutieli, 2001).

Lastly, in comparisons against baseline, per-fROI analyses against empirical null distributions are also reported, which aim to ensure that, at the finest grain (each individual fROI and condition, across participants), differences from baseline are independent of assumptions regarding data normality. These tests were also FDR corrected for multiple comparisons for all fROIs within a network, and across all seven conditions of interest.

## 3. Results

### 3.1. Results of reality check and replication analyses

#### Reality check #1: ISCs in perceptual cortices

We examined ISCs for the conditions of interest in early auditory and visual cortex, grouping the conditions by presentation modality. As expected, we observed stronger ISCs for the auditory conditions in the auditory cortex, and stronger ISCs for the visual conditions in the visual cortex (**Figure 3**). The linear mixed-effects (LME) regression (see ***Section 2.2***) revealed a strong crossover interaction (beta = 0.648, SE = 0.046, *t*(89.79) = 13.970, *p =* 10^-25^).

**Figure 3.**
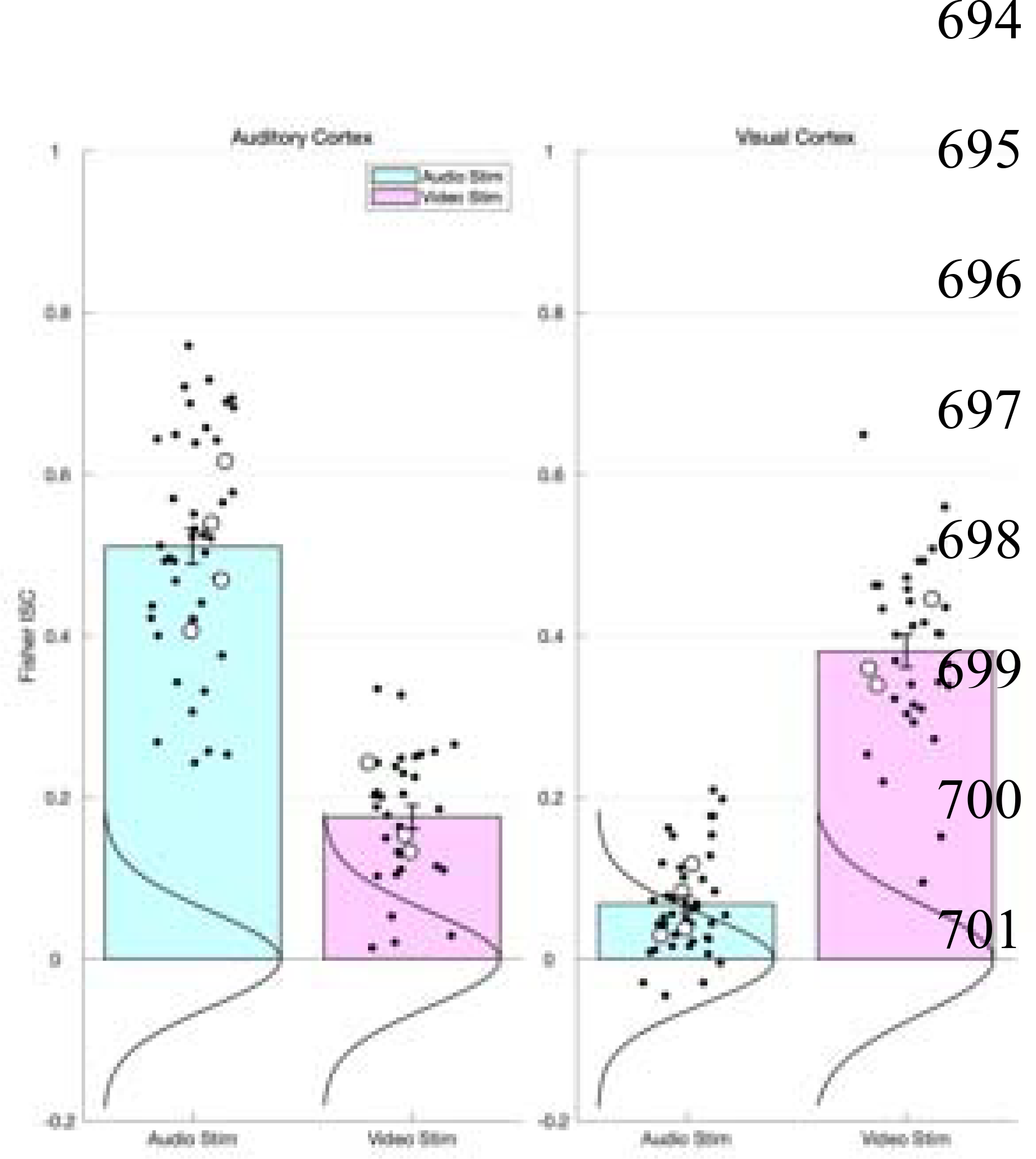
A reality-check analysis showing the expected double dissociation in ISCs in perceptual (visual and auditory) cortices. Bars correspond to Fisher-transformed inter-subject correlation (ISC) coefficients (Pearson’s r) in early visual and auditory cortex to the conditions of interest, grouped by modality of presentation (all +Language conditions: story, audio play, dialog, expository text were auditorily presented; the remaining conditions – animated short film, live action movie clip, and intentional shapes animation – were visually presented). Error bars are standard errors of the mean by participants. Black dots correspond to the individual participants’ values. Large unfilled circles correspond to individual condition averages. Vertical curves are Gaussian fits to empirical null distributions.

**Figure 4.**
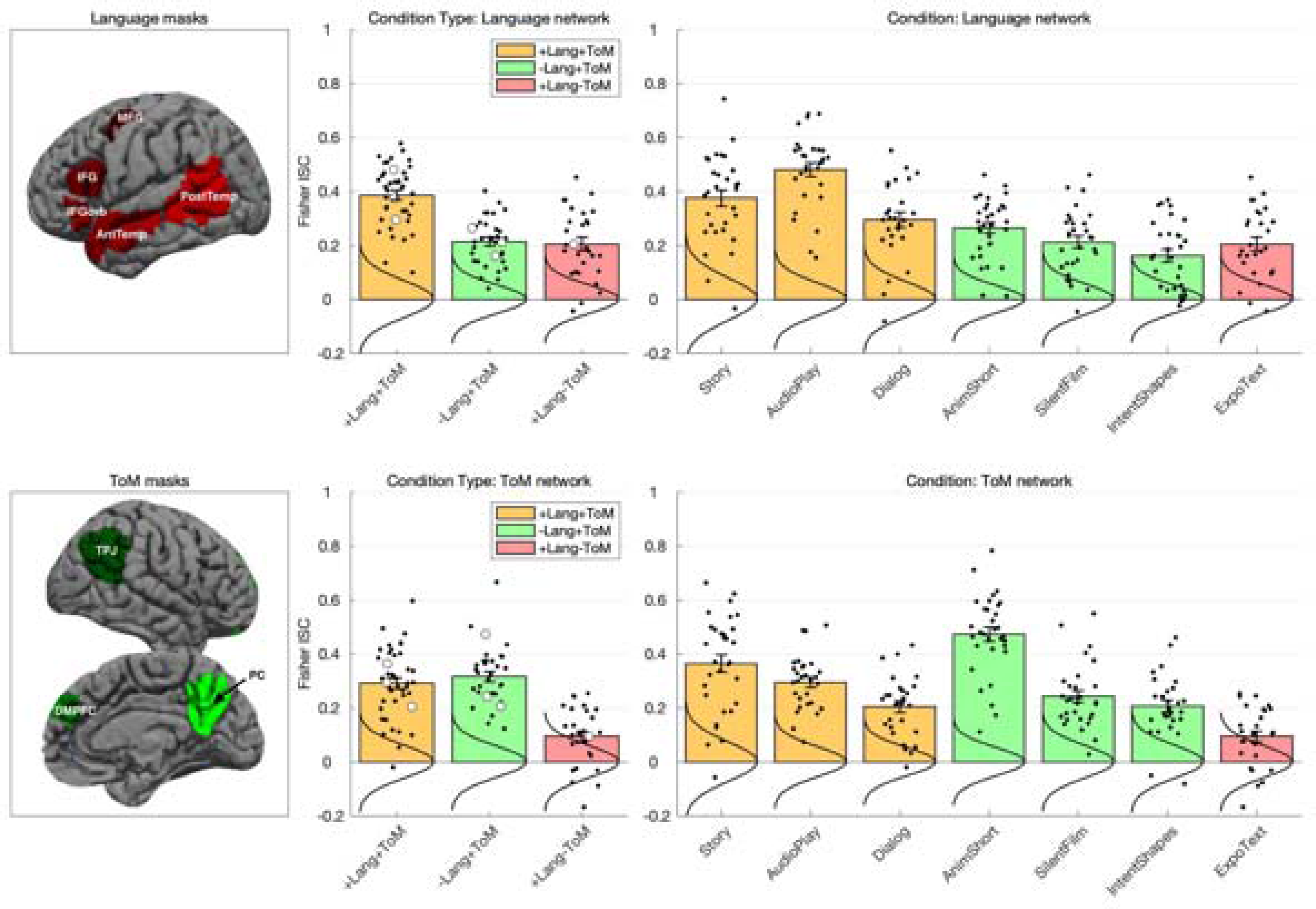
**Left.** Masks within which individual functional regions of interest (fROIs) were defined for each network: Language (Top, red): IFGorb, inferior frontal gyrus, orbital portion; IFG, inferior frontal gyrus; MFG, middle frontal gyrus; AntTemp, anterior temporal cortex; PostTemp, posterior temporal cortex (only the classic, left-hemisphere language regions were included in all analyses). ToM (Bottom, green): TPJ, temporoparietal junction; DMPFC, dorsomedial prefrontal cortex; PC, posterior cingulate cortex and precuneus. Both the right-hemisphere (shown) and left-hemisphere ToM regions were included (six regions total). **Middle.** Average inter-subject correlations (ISCs) per condition type in the language (top) and ToM (bottom) networks. Bars correspond to Fisher-transformed ISC coefficients (Pearson’s r), averaged across regions of interest within each network, separately per condition. Colors represent condition types: +Lang+ToM, orange, -Lang+ToM, green, +Lang-ToM, red. Error bars are standard errors of the mean by participants. Black dots correspond to the individual participants’ values. Large unfilled circles correspond to individual condition averages, shown individually in the right-most panels. Vertical curves are Gaussian fits to empirical null distributions. The key pattern is as follows. The ToM network tracks +ToM materials in both +Linguistic and -Linguistic conditions, but shows weak tracking of the -ToM stimulus. The language network preferentially tracks +Linguistic materials over -Linguistic ones, and it tracks +Linguistic materials in both +ToM and -ToM conditions. **Right.** ISCs per individual naturalistic stimuli (‘conditions’); conventions are as in middle panels. A more detailed characterization of the two networks’ ISC profiles.

Notably, we also observed that the visual areas weakly but reliably tracked the auditory conditions (beta *=* 0.066, SE = 0.012, *t*(38.10) = 5.398, *p* = 10^-7^), and the early auditory areas reliably tracked the visual conditions (beta = 0.175, SE = 0.020, *t*(24.15) = 8.573, *p* = 10^-10^). We return to the interpretation of these effects in the Discussion. For the time being, we note that care must be taken in interpreting deviations of the ISCs during ‘active’ (cf. resting state) conditions from baseline.

#### Reality check #2: Resting state ISCs

To exclude the possibility that the ISCs in the critical analyses are driven by scanner noise or preprocessing/analysis procedures, we measured ISCs across a subset of ten participants who were scanned in a 5-minute resting state condition (Hasson et al., 2004). The ISCs during rest did not significantly differ from baseline in either the language or ToM networks, as assessed with an LME regression, or against the empirical null distribution. This analysis suggests that any above-baseline ISCs for our critical conditions are not an artifact of data acquisition, preprocessing, or analysis procedures.

#### Replication analysis: Closer tracking of linguistic input by language regions than by domain-general Multiple Demand (MD) regions

We successfully replicated the key finding for the narrative condition (beta = 0.181, SE = 0.039, *t*(34.89) = 4.664, *p* = 1^5^; *p* values are FDR-corrected for the four linguistic conditions), and extended it to the audio play (beta = 0.307, SE = 0.052, *t*(34.23) = 5.944, *p* = 10^-7^), the dialog (beta = 0.157, SE = 0.039, *t*(33.05) = 4.014, *p* = 10^-5^), and the expository text (beta = 0.129, SE = 0.036, *t*(36.75) = 3.543, *p* = 10^-4^) conditions (**Supplemental Figure A3**). These results suggest that across diverse kinds of linguistic stimuli, the language network’s activity is more tightly coupled with the inputs, compared to the domain-general MD regions’ activity.

### 3.2. Results of critical analyses

#### Evidence for dissociation between the language and ToM networks in direct network comparisons

We tested three key predictions, which, if supported, provide evidence in favor of language-ToM dissociability in naturalistic settings. First, the ToM network should track conditions rich in ToM content irrespective of whether these conditions are linguistic or non-linguistic, whereas the language network should track linguistic conditions more strongly than non-linguistic ones.

Indeed, we found a network (language, ToM) x condition type (*linguistic*: narrative, audio play, dialog, expository text; *non-linguistic*: animated film, live action film clip, Heider & Simmel-style animation) interaction (beta = 0.191, SE = 0.059, *t*(84.06) = 3.270, *p* = 0.002). We also found main effects of condition type and network: the linguistic conditions were—on average across networks—tracked more strongly than the non-linguistic conditions (beta = 0.124, SE = 0.0459, *t*(96.47) = 2.700, *p* = 0.008), and the language network tracked the conditions more strongly, on average, than the ToM network (beta = 0.096, SE = 0.039, *t*(86.87) = 2.487, *p* = 0.015).

Second, for non-linguistic conditions, the ToM network should exhibit stronger tracking of conditions with mental state content than the language network. Indeed, the ToM network showed higher ISCs than the language network (beta = 0.095, SE = 0.043, *t*(35.59) = 2.197, *p* = 0.035).

And third, for the linguistic condition without mental state content (the expository text), the language network should exhibit stronger tracking than the ToM network. Indeed, the language network showed higher ISCs than the ToM network (beta = 0.111, SE = 0.044*, t*(19.10) = 2.507, *p* = 0.021).

The same qualitative pattern obtains when the non-verbal ToM localizer is used to define the ToM fROIs (**Supplemental Figure C1**).

In this section, we examine more closely the detailed pattern of ISCs in the two networks of interest. The first aim of these analyses is to establish that the observed dissociation is not driven by particular conditions or regions within the networks, but rather, that different aspects of the data provide convergent support for the dissociation. The second aim is to highlight aspects of the ISC pattern that are not consistent with a complete language-ToM dissociation, and thus to evaluate the strength of counter-evidence in favor of the null hypothesis, that language and ToM are not dissociable in naturalistic cognition (see **Supplemental Figure B1** for ISCs per fROI for each network and condition).

#### ToM network

First, the ToM network reliably tracked each of the six conditions with mental state content: story (beta = 0.363, SE = 0.037, *t*(26.55) = 9.693, *p* = 10^-10^); *p* values are FDR-corrected for seven conditions—we are including all conditions in the correction, not only the +ToM conditions), audio play (beta = 0.295, SE = 0.028, *t*(10.54) = 10.716, *p* = 10^-7^), dialog (beta = 0.203, SE = 0.026, *t*(16.40) = 7.797, *p* = 10^-7^), animated short (beta = 0.472, SE = 0.053, *t*(9.35) = 8.830, *p* = 10^-6^), live action film (beta = 0.241, SE = 0.035, *t*(11.18) = 6.869, *p* = 10^-6^), and Heider & Simmel (H&S) style animation (beta = 0.205, SE = 0.026, *t*(15.72) = 7.838, *p* = 10^-7^). Moreover, in tests against the empirical null distributions these effects were significant in every ToM fROI with the exception of the Dialog condition in the RH Posterior Cingulate/Precuneus, (all other *ps* < 0.04, FDR-corrected for the six fROIs and seven conditions).

Second, the ToM network showed no preference for linguistic *vs.* non-linguistic conditions with mental state content (beta = 0.018, SE = 0.043, *t*(49.74) = 0.675, *n.s.*), consistent with these regions’ role in representing mental states irrespective of how this information is conveyed (e.g., Jacoby et al., 2016).

And third, the ToM network tracked the linguistic conditions with mental state content (story, audio play, dialog) more strongly than the one without mental state content (expository text) (beta = 0.199, SE = 0.037, *t*(30.69) = 5.344, *p* = 10^-7^), suggesting that the network represents mental state information in linguistic signals, rather than the linguistic signal itself. However, the ToM network did exhibit weaker but significantly above-baseline tracking of the expository text (beta *=* 0.093, *SE* = 0.022, *t*(22.22) = 4.165, *p* = 0.001). In per-fROI tests against the empirical null distributions, this effect was only reliable in the LTPJ (*p* = 0.015; all other *p*s > 0.05; FDR-corrected for the six fROIs). We consider possible explanations in the Discussion.

#### Language network

First, the language network reliably tracked each of the three linguistic conditions with mental state content: story (beta = 0.374, SE = 0.041, *t*(13.27) = 9.171, *p* = 10^-7^; *p* values are FDR-corrected for seven conditions), audio play (beta = 0.479, *SE* = 0.046, *t*(8.75) = 10.370, *p* = 10^-6^), and dialog (*r* = 0.294, *SE* = 0.045, *t*(9.72) = 6.491, *p* = 10^-4^). Moreover, these effects were significant in every language fROI, in tests against the empirical null distributions (*p*s < 0.01, FDR-corrected for the five fROIs and seven conditions).

Second, importantly, the language network also reliably tracked the linguistic condition with no mental state content (beta = 0.204, *SE* = 0.044, *t*(8.72) = 4.668, *p* = 10^-4^), and this effect, too, was significant in every language fROI (*ps* < 0.03, FDR-corrected for the five fROIs). This result suggests that mental state content is not necessary to elicit reliable ISCs in the language network.

And third, the language network showed stronger tracking of linguistic relative to non-linguistic conditions with mental state content, (beta = 0.177, SE = 0.043, *t*(43.54) = 4.155, *p* = 10^-5^). This result suggests a special role for linguistic input in driving the network’s responses.

However, the language network exhibited some patterns that might be taken to suggest that mental state content—or social information more generally—is, to some extent, important for linguistic processing. First, the language network tracked the linguistic conditions with mental state content more strongly than the linguistic condition with no mental state content, i.e., the expository text (beta = 0.188, SE = 0.061, *t*(25.67) = 3.050, *p* = 0.005). This result may be taken to suggest that mental state content contributes to the language network’s input tracking over and above the linguistic content alone. This interpretation warrants caution, however. In particular, reflecting the general challenges of naturalistic stimuli (see Discussion), the linguistic condition with no mental state content is not matched to the linguistic conditions with mental state content on various potentially relevant features, from how engaging they are, which could influence the depth of linguistic encoding, to specifically linguistic properties (e.g., lexical and syntactic complexity), which could also affect the strength of ISCs (e.g., Shain, Blank et al., 2020; Wehbe et al., 2021). Furthermore, only a single linguistic condition with no mental state content was included in the current study, making it difficult to rule out idiosyncratic features driving the difference.

And second, the above-baseline ISCs in the language network for the non-linguistic conditions— although weaker than those for the linguistic conditions—are also notable, suggesting some degree of reliable tracking in the language network for non-linguistic meaningful information (see also Ivanova et al., 2020 for evidence of reliable responses in the language network to visual events).

## 4. Discussion

Much prior work in cognitive neuroscience has suggested—based on traditional controlled experimental paradigms—that the network of brain regions that support linguistic interpretation and the brain regions that support mental state reasoning are distinct (e.g., Saxe & Kanwisher, 2003; Mason & Just, 2009; Fedorenko et al., 2011; Mar, 2011; Deen et al., 2015; Paunov, 2019). However, such paradigms differ drastically from real-world cognition, where we process rich and complex information. And linguistic and social cognition seem to be strongly intertwined in everyday life. Here, we tested whether the language and Theory of Mind networks are dissociated in their functional profiles as assessed using the inter-subject correlation (ISC) approach, where neural activity patterns are correlated across individuals during the processing of naturalistic materials (e.g., Hasson et al., 2004, 2008). Following Blank & Fedorenko (2017), we combined the ISC approach with the power of individual-participant functional localization (e.g., Brett et al., 2002; Saxe et al., 2006; Fedorenko et al., 2010; Nieto-Castañón & Fedorenko, 2012). This synergistic combination has two key advantages over the whole-brain voxel-wise ISC approach, where individual brains are first anatomically aligned and, then, each stereotaxic location serves as a basis for comparing signal time courses across participants. First, relating the resulting cortical topography of ISCs to the topography of known functional brain networks can only proceed only through “reverse inference” based on anatomy (Poldrack, 2006; Fedorenko, 2021). Instead, evaluating signal time courses from functionally defined regions ensures interpretability, and allows us to straightforwardly link our findings to the wealth of prior studies characterizing the response profiles of our two networks of interest. And second, this approach allows us to directly test the correlations in the language network against those in the ToM network. Such an explicit comparison between networks allows for stronger inferences compared with those licensed when each network is separately tested against a null baseline and differences across networks are indirectly inferred (e.g., see Nieuwenhuis et al., 2011, for discussion).

We examined the ISCs in the language and ToM networks during the processing of seven naturalistic conditions: three linguistic conditions with mental state content (+linguistic, +ToM), three non-linguistic conditions (silent animations and live action films) with social content but no language (-linguistic, +ToM), and a linguistic condition with no social content (+linguistic, -ToM). We found reliable differences in the ISC patterns between the language and ToM networks, in support of the hypothesis that language and Theory of Mind are dissociable even during the processing of rich and complex naturalistic materials. In particular, the ToM network tracked materials rich in mental state content irrespective of whether this content was presented linguistically or non-linguistically (see also Jacoby et al., 2016), but it showed only weak tracking of the stimulus with no mental state content. In contrast, the language network preferentially tracked linguistic materials over non-linguistic ones, and it did so regardless of whether these materials contained information about mental states.

These results expand on the existing body of knowledge about language and social cognition, with both theoretical and methodological implications. Critically, the observed dissociation extends prior findings of dissociable functional profiles between the language and the ToM networks during task-based paradigms to rich naturalistic conditions. This result suggests that the two networks represent *different kinds of information*. (They may also perform *distinct computations* on the perceptual inputs, though the idea of a canonical computation carried out across the cortex is gaining ground (e.g., Keller & Mrsic-Flogel, 2018; Fedorenko & Shain, in press), and predictive processing seems like one likely candidate (e.g., Koster-Hale & Saxe, 2013; Shain, Blank et al., 2020).) In particular, the language regions appear to track linguistic features of the input (see also Shain, Blank et al., 2020; Shain et al., 2021; Wehbe et al., 2021).

Our results extend prior findings from ISC paradigms (Wilson et al., 2008; Lerner et al., 2011; Honey et al., 2012; Regev et al., 2013; Silbert et al., 2014; Schm□lzle et al., 2015; Blank & Fedorenko, 2017), which all used materials rich in mental state content, as is typical of linguistic information, to a stimulus that is largely devoid of information about mental states—an expository text. Strong tracking of the latter stimulus aligns with prior findings from task-based paradigms of robust responses to linguistic materials with little or no mental state content (e.g., Deen et al., 2015; Jacoby & Fedorenko, 2020).

The ToM regions, in contrast, appear to track some features related to representing mental states across diverse kinds of representations (linguistic materials, animations, including highly abstract and minimalistic ones, and live action movies), again aligning with prior findings from task-based paradigms (e.g., Fletcher et al., 1995; Gallagher et al., 2000; Castelli et al., 2000; Vogeley et al., 2001; Ruby & Decety, 2003; Saxe & Kanwisher, 2003; Saxe, Schulz, & Jiang, 2006; Jacoby et al., 2016). It is important to keep in mind that the fact that the two networks are dissociable does not imply that they do not interact. Indeed, Paunov et al. (2019) reported reliably above chance correlations in the patterns of inter-regional synchronization between the language and ToM networks, suggesting some degree of functional integration.

On the methodological level, these results vindicate the divide-and-conquer strategy in general, and the functional localization approach (e.g., Brett et al., 2002; Saxe et al., 2006; Fedorenko et al., 2010) in particular. Language and Theory of Mind appear to be distinct, supported by dissociable cortical networks for the processing of linguistic vs. mental state information, at least in adulthood (see also Braga et al., 2020). It is therefore justifiable to study each cognitive faculty and each network separately, although further probing the mechanisms of their potential interactions is equally important.

Although the overall pattern clearly supports a language-ToM dissociation between the two networks, some aspects of the results are not in line with a complete dissociation. In particular, i) the language regions show reliable tracking of non-linguistic conditions with mental state content; ii) the language regions show stronger tracking of linguistic conditions with than without mental state content; and iii) the ToM regions show weak but reliable tracking of the linguistic condition with no mental state content. These findings may be due to methodological limitations: the above-baseline ISCs (especially the relatively weak ones) may not reflect stimulus tracking. Although we have ruled out the possibility that the above-baseline ISCs are driven by acquisition, preprocessing, or analysis artifacts in our reality-check analysis of resting state data, they could be driven by other, non-mutually-exclusive, factors. One possibility is that inter-network interactions could induce ISCs. In particular, given that the language and the ToM network show some degree of synchronization in activity during naturalistic cognition (Paunov et al., 2019), the ToM network’s tracking of a linguistic stimulus with no mental state content, for example, may be due do the fact that the language system is tracking this stimulus, and there is some ‘leakage’ of this tracking to the ToM network through inter-network synchronization. Similarly, the language network’s preference of linguistic stimuli with mental state content over those without mental state content may be due to the leakage from the ToM network.

Another possibility is that the incomplete dissociation between language and Theory of Mind (at least with respect to (i) and (iii) above) may be at least partly attributable to pragmatic processing, arguably present across all ‘conditions’: the non-linguistic ToM conditions are still ‘story-like’ and hence communicative and the linguistic non-ToM stimulus arguably still requires attribution of communicative intentions, as discussed in the introduction. The division of labor between the language and ToM networks in pragmatic inference is an exciting future direction. Demonstrating that these two networks are, in the first place, dissociable, even during rich naturalistic cognition––the goal of the present study––is an important step to pursuing this line of research. With this groundwork in place, neuroimaging evidence can be increasingly brought to bear on the question of which aspects of pragmatics require mental state reasoning, as evidenced by the engagement of the ToM network. Given the broad scope of pragmatics, encompassing diverse heterogeneous phenomena, an empirically motivated pragmatic taxonomy may be developed by investigating whether some classes of pragmatic inference are resolved within the language network proper (e.g., ‘lower-level’ inferences about lexical or syntactic ambiguity) whereas others (e.g., establishing discourse coherence, or understanding irony) require the ToM network.

Finally, the stimuli themselves may be too confounded to fully dissociate the respective contributions of language- and ToM-related components. For example, in linguistic stimuli, mental state attribution often requires particular syntactic structures—sentential complements.

More generally, the use of naturalistic materials, despite its advantages (e.g., Hasson et al., 2018; Sonsukare et al., 2019; Nastase et al., 2020), is associated with a host of challenges. The key one, mentioned above, is that certain features are necessarily confounded in naturalistic settings, and can only be dissociated through careful experimentation and altering the natural statistics of the input. Relying on naturalistic materials alone can lead to wrong conclusions about the cognitive and neural architecture. This problem is especially pronounced in studying the relationship between language and social cognition given that language is primarily used in social settings and to share socially relevant information. The use of linguistic materials with no social information, and of non-linguistic mental-state-rich materials has been critical, here and in earlier studies, to uncover the dissociation that holds between the language and ToM systems. Another challenge associated with the use of naturalistic materials is that they are difficult, or altogether impossible, to match for diverse properties bound to affect neural responses. Again, this problem presents a particular challenge in comparing responses to materials rich in social information vs. devoid of such information given that the former are, almost by definition, going to be more engaging and exciting given the social nature of primates, including humans (e.g., Aronson, 1980; Cheney & Seyfarth, 1990; Tomasello, 2014). Possible ways to address these concerns could involve a) characterizing the natural statistics of co-occurrences between linguistic and social processing in order to better understand how well the naturalistic stimuli reflect those statistics and perhaps altering naturalistic conditions to allow dissociating features that commonly co-occur in life; b) developing novel neural analysis methods to isolate the components of neural signals attributable to a particular cognitive process / brain network (e.g., using analytic methods well-suited to high-dimensional data such as independent components analysis across both cortical space and large feature spaces representative of naturalistic environments (e.g., Norman-Haignere et al., 2015), or across both cortical space and time (e.g., probabilistic ICA, PICA; see Beckmann et al., 2005 for overview); and c) carefully annotating naturalistic materials and performing reverse correlation analyses (e.g., Hasson et al., 2004; Richardson et al., 2018) in an effort to understand the precise features that elicit increases in neural responses in different brain regions. The latter approach may be particularly informative with respect to the question of possible *gradation* of demands on language and ToM processing both within naturalistic stimuli and across ‘conditions’ that we here grouped in the same ‘type’ or general categories of +/-ToM and +/-Language. Our results seem to suggest that considerable within-category heterogeneity exists in the degree of stimulus tracking (e.g., in the language network, the dialog is tracked less strongly than the narrative, and in the ToM network, the Heider & Simmel-style animation is tracked less strongly than the animated film. Our dataset is not ideally suited for such investigation because it includes only a single instance of each ‘condition’, and no attempt was made to match the conditions on at least *some* dimensions that may improve homogeneity, but this is a promising direction for future work.

A few other exciting future directions are worth highlighting, some of which may build directly on the approaches introduced and the findings reported in the current study. First, the language and ToM networks appear to be dissociable in the adult mind and brain. However, it is possible—perhaps even plausible—that this dissociation emerges over the course of development. Prior neuroimaging work has shown that the ToM network becomes gradually more specialized for mental state attribution (e.g., Saxe, 2009; Gweon et al., 2012; Richardson et al., 2018), and this specialization appears to be protracted with delayed language acquisition (Richardson et al., 2020). Very little is known about how specialization for linguistic processing (e.g., Fedorenko et al., 2011; Monti et al., 2012) emerges. Perhaps early on in development, a set of lateral frontal, lateral temporal, and midline cortical areas are tuned to *any* socially relevant information, and these areas later fractionate into those specialized for processing linguistic signals vs. those for mental state attribution vs. those that support many other kinds of social signals, both visual (e.g., eye gaze, facial expressions, gestures) and auditory (e.g., non-verbal vocalizations, prosodic information, speech acoustics). This fractionation is likely driven by computational and metabolic advantages of localized processing (e.g., Barlow, 1995; Foldiak & Young, 1995; Chklovskii & Koulakov, 2004; Olshausen & Field, 2004; see Kanwisher, 2010, for discussion). Probing linguistic and social cognition across the lifespan will be critical to understand how the two networks form and develop, leading to the segregation we observe in the adult brain.

Second, as noted above, given the likely frequent interactions between the language and ToM networks (including their most strongly dissociated components), searching for possible mechanisms of those interactions (e.g., Paunov et al., 2019) seems critical. This would require a combination of studies characterizing the patterns of anatomical connections for the language and ToM regions (e.g., Saur et al., 2010; Wiesmann et al., 2017) and studies probing online interactions using methods with high temporal resolution, like MEG or intracranial recordings.

Third, we are still a long way away from a mechanistic-level understanding of what the language or the ToM regions do. The use of naturalistic stimuli, including in the context of the ISC approach, is promising. In particular, by examining the points in the stimulus where most participants show increases in neural activity can help generate (and subsequently test) specific hypotheses about the necessary and sufficient features of the input that are required to elicit neural responses in the relevant brain regions.

To conclude, we have demonstrated that the dissociation between the language and ToM networks that has been previously reported based on traditional task paradigms, robustly generalizes to rich naturalistic conditions. However, the precise nature of each network’s representations and computations, the emergence of these networks in development, and the mechanisms for information sharing between them remain to be discovered.

## Supporting information

Supplementary Results

## Acknowledgments

We would like to acknowledge the Athinoula A. Martinos Imaging Center at the McGovern Institute for Brain Research at MIT, and its support team (Steve Shannon and Atsushi Takahashi). We thank former and current EvLab members (especially Brianna Pritchett and Caitlyn Hoeflin) for their help with data collection, Amaya Arcelus for help with creating the Heider & Simmel style animation, Caroline Arnold for help with recording the dialog condition, and Jessica Chen and Yotaro Sueoka for help with manuscript preparation. This research and EF were supported by R01 awards DC016607 and DC016950 from NIDCD, a grant from the Simons Foundation to the Simons Center for the Social Brain at MIT, and the Newton Brain Science Award from the Brain and Cognitive Sciences Department at MIT.

## Contributions

AP, IB and EF developed the concept for the study. AP, IB and JG selected / created experimental materials. AP, IB, OJ and ZM collected and analyzed the data. AP performed the final version of the statistical analyses for the current study that are reported here. AP, IB, and EF wrote the manuscript with input from other co-authors.

