## Supplementary Results for "Differential tracking of linguistic vs. mental state content in naturalistic stimuli by language and Theory of Mind (ToM) brain networks"

**Section A.** *Replication analysis: Closer tracking of linguistic input by language regions than by domain-general Multiple Demand (MD) regions.*

***SI Figure A1.*** *Paradigm used to localize the MD network.* Harder and easier versions of a spatial working memory task (location memory) were contrasted, each followed by a two-alternative forced choice question and feedback.

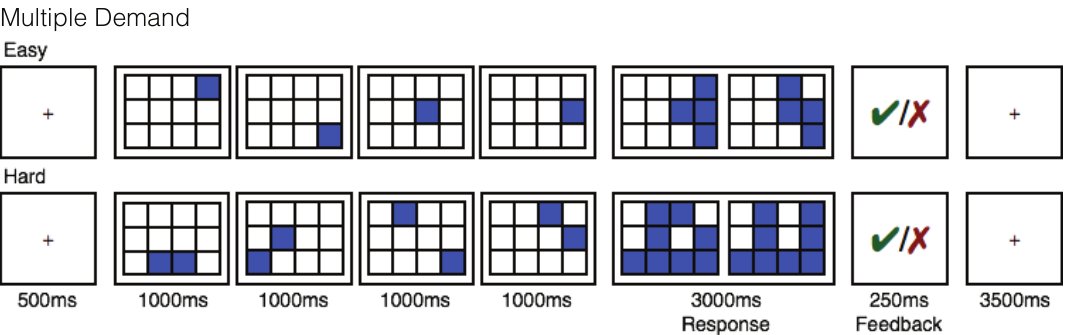

***SI Figure A2.*** *MD network masks.* Nine anatomically defined MD masks were used in each hemisphere (Tzourio-Mazoyer et al., 2002; only the left-hemisphere ones shown here), including regions in the middle frontal gyrus *(MFG)* and its orbital part *(MFGorb)*, the opercular part of the inferior frontal gyrus (*IFGop)*, the precentral gyrus *(PrecG),* the superior *(ParSup)* and inferior *(ParInf)* parts of the parietal lobe, the insula *(Insula),* the supplementary motor area *(SMA)*, and the cingulate cortex *(ACC).*

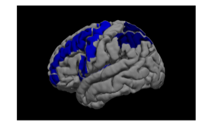

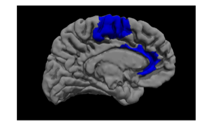

***SI Figure A3.*** *Replication of the language-MD dissociation in narrative processing (Blank & Fedorenko, 2017) and extension to other linguistic conditions.* All language vs. MD effects are significant in LME regressions with random intercepts per fROI and participant after FDR correction (see Main Methods & Results). Error bars are standard errors of the mean by participants. Black dots correspond to the individual participants’ values. Vertical curves are Gaussian fits to empirical null distributions.

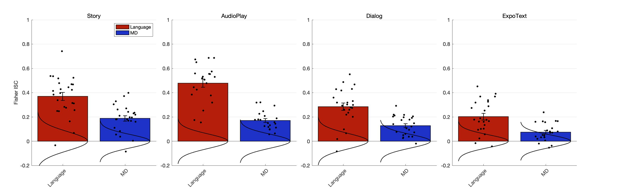

**Section B.** *Inter-subject correlations per functional region of interest (fROI).*

***SI Figure B1.*** *Individual fROI ISCs in the language (A) and ToM (B) networks for each of the seven critical conditions*. The left panel in each plot shows the average ISC across ‘conditions’ of a given type, shown individually on the right. Color codes correspond to condition types: +Language+ToM (yellow), -Language+ToM (green), and +Language-ToM (red). Error bars are standard errors of the mean by participants. Black dots correspond to the individual participants’ values. Vertical curves are Gaussian fits to empirical null distributions.

A. *Language fROIs*

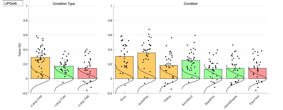

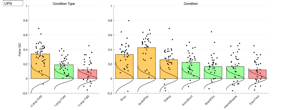

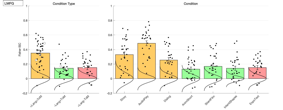

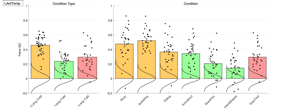

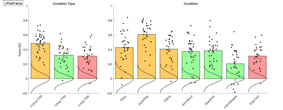

B. *Theory of Mind fROIs*

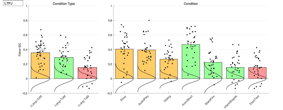

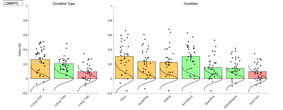

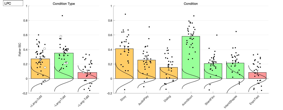

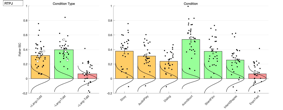

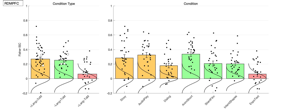

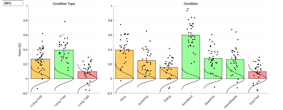

**Section C.** *Inter-subject correlations with a non-verbal Theory of Mind contrast*.

To examine whether the network dissociation holds with ToM regions localized with a non-verbal task, we defined ToM fROIs with the non-verbal AnimShort condition (-Lang+ToM) and reproduced Figure 3 in the manuscript.

Specifically, the AnimShort stimulus has been previously (Jacoby et al., 2016) coded into segments high in mental state content (‘mental’; e.g., character falsely believes they have been abandoned by a companion; 4 events, 44 sec total) and segments high in physical information (‘physical’; e.g., wide shot of clouds, storks flying; 3 events, 22 sec total). In the subset of subjects who were scanned on this condition (n=34), we defined ToM fROIs with this non-verbal mental > physical contrast and extracted time series for all conditions of interest from these fROIs. The rest of the procedure for obtaining ISCs is as described in the text.

Qualitatively, the same pattern of ISCs across condition types obtains in the ToM network (see Figure 4 in the main text for comparison), with slightly reduced ISCs for the +Language conditions, hinting toward a stronger language-ToM dissociation when a non-verbal ToM contrast is used, consistent with naturalistically dissociable language and ToM networks.

Note that the correlation for the AnimShort stimulus is obtained with the same data as used to define the ToM fROIs. Given that the voxels are pre-selected to maximize the difference between mental and physical segments of the stimulus, the increased ISC for the AnimShort condition cannot be interpreted due to non-independence.

***SI Figure C1.*** *ISCs in the language (top) and Theory of Mind (bottom) networks, with ToM fROIs derived using a non-verbal ToM contrast.* **Left.** Masks within which individual functional regions of interest (fROIs) were defined for each network: Language (Top, red): IFGorb, inferior frontal gyrus, orbital portion; IFG, inferior frontal gyrus; MFG, middle frontal gyrus; AntTemp, anterior temporal cortex; PostTemp, posterior temporal cortex (only the classic, left-hemisphere language regions were included in all analyses). ToM (Bottom, green): TPJ, temporoparietal junction; DMPFC, dorsomedial prefrontal cortex; PC, posterior cingulate cortex and precuneus. Both the right-hemisphere (shown) and left-hemisphere ToM regions were included (six regions total). **Middle.** Average inter-subject correlations (ISCs) per condition type in the language (top) and ToM (bottom) networks. Bars correspond to Fisher-transformed ISC coefficients (Pearson’s r), averaged across regions of interest within each network, separately per condition. Colors represent condition types: +Lang+ToM, orange, -Lang+ToM, green, +Lang-ToM, red. Error bars are standard errors of the mean by participants. Black dots correspond to the individual participants’ values. Large unfilled circles correspond to individual condition averages, shown individually in the right-most panels. Vertical curves are Gaussian fits to empirical null distributions. The key pattern is qualitatively the same as that reported in Figure 3, with a verbal ToM localizer, as follows. The ToM network tracks +ToM materials in both +Linguistic and -Linguistic conditions, but shows weak tracking of the -ToM stimulus. The language network preferentially tracks +Linguistic materials over -Linguistic ones, and it tracks +Linguistic materials in both +ToM and -ToM conditions. **Right.** ISCs per individual naturalistic stimuli (‘conditions’); conventions are as in middle panels.

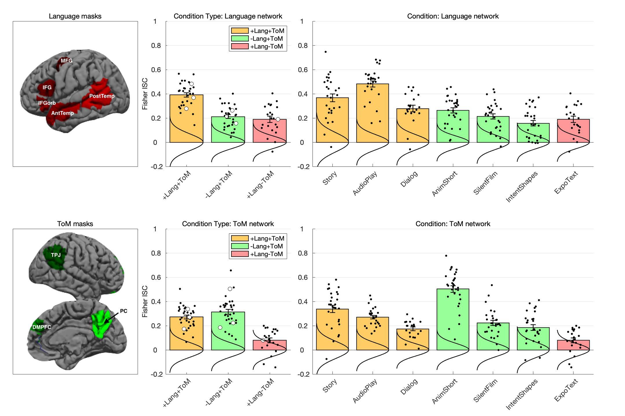
